# Functional Profiling of 2,193 *ASS1* Missense Variants: Insights into Variant Pathogenicity and Epistatic Interactions in Citrullinemia Type I

**DOI:** 10.1101/2025.09.17.676623

**Authors:** Russell S. Lo, Gareth A. Cromie, Michelle Tang, Amy Sirr, Ljubica Caldovic, Hiroki Morizono, Nicholas Ah Mew, Andrea Gropman, Aimée M. Dudley

## Abstract

Sequence variants in the urea cycle gene argininosuccinate synthase (*ASS1*) cause Citrullinemia type 1 (CTLN1), a rare autosomal recessive disease. Mechanistically, reduction in argininosuccinate synthetase (ASS) enzyme activity impairs the urea cycle, leading to an accumulation of citrulline and neurotoxic ammonia. Disease severity varies according to the degree of enzyme impairment, ranging from severe neonatal forms (classic citrullinemia) to milder, late-onset forms that may manifest in childhood or adulthood. We established a high-throughput yeast functional assay of human ASS and individually measured the impact of 2193 amino acid substitutions, representing 90% of all single nucleotide variant (SNV)-accessible substitutions. When benchmarked against existing clinical variant annotation, our assay distinguishes known benign variants from strong loss of function pathogenic variants, enabling identification of a functional score threshold below which variants show clinically relevant impairment of ASS activity. Using the ACMG OddsPath framework, our assay meets PS3_supporting criteria for pathogenicity classification and achieves full PS3-level strength when variants observed as homozygotes in other primates are used as benign proxies for calibration. These results provide direct functional evidence to inform reclassification of *ASS1* missense variants, supporting their clinical interpretation and diagnostic utility. Mapping functional scores onto the protein structure, we confirmed that residues involved in catalysis are highly sensitive to substitution. In addition, we identified residues from adjacent subunits of the ASS homotetramer that form compound active sites. Assaying these positions revealed a capacity for intragenic complementation consistent with a variant sequestration model: a form of positive epistasis in which deleterious variants from different subunits are sequestered into only a subset of active sites, restoring function in the remaining variant-free sites. The discovery of intragenic complementation in ASS reveals a novel mode of functional interaction with clinical implications for interpreting variant combinations in heterozygous individuals.

## Background

ASS1 encodes human argininosuccinate synthetase (ASS), one of the enzymes of the urea cycle, a pathway that converts ammonia, a toxic waste product of protein metabolism, into urea. Specifically, ASS catalyzes the formation of argininosuccinate from citrulline and aspartic acid. Deleterious sequence variants in ASS1 cause the rare autosomal recessive disease citrullinemia type I (CTLN1). CTLN1 is the third-most common of the urea cycle disorders, with symptoms driven by the accumulation of citrulline and bouts of hyperammonemia, in which the brain is exposed to toxic levels of ammonia, leading to neurological damage.

The severity of citrullinemia type I (CTLN1) correlates with the level of residual ASS enzyme activity. Absent or negligible ASS function result in life-threatening neonatal disease onset. This classic form of citrullinemia typically presents within the first few days of life, with symptoms including lethargy, poor feeding, failure to thrive, vomiting, seizures, coma, and, if untreated, death. Partial deficits in ASS activity may result in a milder, late-onset form of the disease, or even a clinically asymptomatic presentation marked only by elevated plasma citrulline and low arginine. However, catabolic stress, such as during the post-partum period or following surgery, infection, fever, or physical trauma, can trigger acute metabolic decompensation events even in individuals with only partial ASS deficiency. These episodes may cause neurological impairment, including abnormal behaviors, seizures, coma, and even death [1]. Thus, even late-onset events can be life-threatening, and recovery may be accompanied by lasting intellectual disabilities.

All 50 U.S. states screen for CTLN1 as part of their newborn screening programs. CTLN1 is included as a core condition on the Recommended Uniform Screening Panel (RUSP), a list of disorders recommended by the Secretary of Health and Human Services (HHS) for state screening programs based on the availability of effective treatments and the demonstrated benefits of early detection. Screening is performed using tandem mass spectroscopy to detect elevated levels of citrulline. If a newborn is positive on both initial and follow-up testing, medical intervention utilizing dietary management with low protein formulas, and ammonia-scavenging drugs can be employed. In more severe cases, liver transplants are considered.

Elevated citrulline levels identified through newborn screening typically prompt follow-up sequencing of the *ASS1* gene to detect pathogenic variants underlying enzyme deficiency. Variant interpretation is guided by the ACMG/AMP framework, which integrates evidence from computational predictions, population frequency, segregation, clinical phenotype, and functional assays [2]. However, because ASS1 deficiency is rare, many variants identified in affected individuals lack sufficient supporting data and are classified as variants of uncertain significance (VUS) [2]. These VUS are not clinically actionable, limiting the diagnostic utility of sequencing-based approaches. As sequencing expands, the number of novel *ASS1* variants with unclear clinical significance is expected to grow. Functional assays have the potential to provide strong orthogonal evidence for both pathogenic and benign classifications. Here, we describe a high-throughput functional assay of ASS enzyme activity characterizing the impact 2193 amino acid substitutions, representing 90% of all single nucleotide variant (SNV)-accessible substitutions in *ASS1*. These results provide additional evidence toward a pathogenic classification for 547 amino acid substitutions in ASS1, including 25 that correspond to ClinVar variants currently classified as VUS.

## Methods

### Plasmid and strain construction

The *S. cerevisiae* strains used in this study (Table S1) were derived from the isogenic lab strain FY4 and grown in rich YPD medium (1% yeast extract, 2% peptone, and 2% glucose) or minimal (SD) medium (without amino acids, 2% glucose), with standard methods for yeast genetic manipulation [3].

To construct yeast strains harboring *ASS1* variants, we first deleted the *ARG1* open reading frame and terminator from FY4 and replaced it with a selectable kanamycin resistance gene and the *S. cerevisiae ADH1* terminator from pFA6a-K*anNT2*. Therefore, in this strain, expression of the kanamycin resistance gene was under control of the *S. cerevisiae ARG1* promoter and the *S. cerevisiae ADH1* terminator.

We optimized the human *ASS1* coding sequence (GenBank: NM_054012.4) for expression in yeast (hereafter *yASS1*, GenBank: PQ374833) by using a custom, in-house method designed to match the codon usage frequency of *S. cerevisiae*. At positions where amino acids are conserved between the yeast and human proteins, the yeast (S288c reference) codon was used. In non-conserved locations, codons encoding the appropriate ASS amino acid were chosen to be as similar as possible to the usage frequency of the codons at corresponding positions in *ARG1*.

Plasmids containing human *ASS1* (*hASS1)* or yeast codon-optimized *ASS1* (*yASS1)* were constructed as follows. Two DNA fragments, one containing both the *ARG1* promoter (*pARG1*) and wild type *ASS1* (*hASS1* or *yASS1* ORFs synthesized by IDT), and one containing the *ARG1* terminator (*tARG1*), the *NatMX* drug selection cassette, and sequence downstream of the *ARG1* terminator (amplified via PCR), were cloned into pUC19 vector with the NEB HiFi DNA Assembly Cloning Kit (New England Biolabs) to create AB596_hASS1 & AB598_yASS1 plasmids (GenBank: PV290047 & PQ374833). The AB598_yASS1 plasmid was used as a template for variant library construction.

### Construction of variant library

We designed the variant library to capture the amino acid substitutions resulting from all SNV-accessible missense sequence variants, excluding the start and stop codons, in the human *ASS1* cDNA sequence (GenBank: NM_054012.4). These SNV-accessible amino acid substitutions were defined relative to the native human *ASS1* cDNA sequence, not the *yASS1* codon-optimized sequence. The nine possible SNVs at each native codon produce 4–7 unique amino acid substitutions. *yASS1* derivatives encoding the complete set of these unique amino acid substitutions (n = 2450) were synthesized (Twist Biosciences) as part of a *pARG1-yASS1-tARG1-NatMX* cassette. This resulted in a variant library in which each well of a 96-well plate contained an approximately equal pool of genotypes encoding each of the 4–7 amino acid substitutions at a given amino acid position.

Separate PCR amplification reactions and subsequent integrative yeast transformations were then performed for each Twist well, i.e., a separate transformation for each amino acid position tested (n = 411). Approximately 300 ng of each PCR-amplified amino acid variant pool was transformed into a MATa haploid *arg1* deletion strain (YAD1268) using standard methods. From each transformation, single colonies were isolated such that a total of 7,704 individual transformants were arrayed into 96-well plates containing rich medium. This number of colonies was chosen such that approximately three to four independent transformants of each variant would be isolated and assayed. For downstream phenotype normalization, each library plate also contained replicates of the following control strains: two deletion (*arg1*Δ*0*) and four wild type (*yASS1*) strains. Stocks were maintained as individual strains in 96-well format.

### Variant library sequence confirmation

In our strain construction pipeline, for each transformant, the target codon harboring the amino acid substitution is known, but the specific variant is not. To determine this, we used a custom MinION (Oxford Nanopore Technologies) long-read sequencing pipeline to sequence the entire open reading frame (as described in [4]). Briefly, individual transformants were pooled in groups of 14 or 15, such that no target codon position was represented more than once in a single pool. These pools and their target codons are described in Supplemental Table S2. Each pool was then sequenced using Oxford Nanopore Flongle flow cells (R9.4.1), and the associated sequences are downloadable from the Sequence Read Archive under accession PRJEB91359 [5]. At each target codon in each pool, the most frequent potential variant codon was identified (candidate variant) as well as the second most frequent variant (second variant), from the set of encodings used by Twist Bioscience for yeast (Supplemental Table S3), using a script as described in [4] (Supplementary Materials, Method S2). Because we know which target codon corresponds to which transformant, this allows us to associate each candidate variant with a single transformant. Any potential secondary mutations were also identified.

The frequency and pattern of MinION sequencing errors is highly variable and depends on the context of surrounding bases. These errors can occur at high frequencies, potentially generating spurious matches to variant codons. However, the error patterns are also reproducible per given DNA sequence, allowing us to develop an error model for each variant, describing the frequency with which it is generated by sequencing errors. This frequency can be compared to the frequency observed for each candidate variant in the pooled sequencing, allowing true variants to be distinguished from sequencing noise.

These data were used to carry out quality control on the variant calls. The candidate variant was provisionally accepted if supported by >=15 reads and if the ratio of first variant codon counts to reference codon counts was >0.02 and <0.2, based on the multiplexing expectation of 1/14 or 1/15 along with some sampling variation. The observed frequency of each candidate variant codon was then compared to the frequency expected under the variant-specific error model. Candidate variants were rejected if their enrichment ratio (observed frequency / error frequency) was <3.3. We also rejected candidate variants if their enrichment ratio was <3.3 times that of the second most frequent variant, or if the enrichment ratio of the second most frequent variant was very high (>10). Finally, candidates were also flagged for later removal if any missense or nonsense secondary mutations, or insertions or deletions (indels), were present in the *yASS1* sequence. The end result of this process was that each transformant was assigned either a high confidence call or was flagged for removal from the analysis.

### Measuring the growth of individual isolates

To prepare for phenotyping, each isolate was picked and grown in rich medium supplemented with the selectable drug and then pinned onto solid rich medium containing 3% glycerol as the carbon source (YPG), to select against cells that were slow growing as a result of loss of respiratory function. Cells were then pinned from YPG solid medium back into non-selective rich liquid medium (YPD) and grown to saturation overnight at 30° C. Next, cultures were pinned onto solid omniplates containing minimal (SD) medium (lacking arginine) with a Biomek i7 robot outfitted with a V&P Scientific 96 floating-pin replicator head. Pinning onto SD plates was done in triplicate, followed by growth at 30° C for 72 h. Plates were photographed with a mounted Canon PowerShot SX10 IS compact digital camera under consistent lighting, camera to subject distance, and zoom. Images (ISO200, f4.5, 1/40 second exposure) were acquired as jpg files. Growth phenotypes were extracted from the images via a custom pipeline, PyPL8 [6]. Briefly, a pseudo patch ‘‘volume’’ for each pinned isolate was generated, consisting of the product (‘‘IntDen’’) of the patch area (obtained via thresholding with Otsu’s method or circle detection) and the mean patch pixel intensity (grayscale) (as described in the Supplemental Methods in [4].

### Data Processing

Normalization was carried out to account for plate variation, relative growth of neighboring patches, and edge effects as described in [4] using a custom script (Supplementary Materials, Method S2). Isolates with secondary mutations or discordant growth estimates were removed. Finally, relative growth of each ASL variant was estimated using a weighted least-squares regression.

### Additional Variants

To maximize the representation of PrimateAI-3D homozygotes, gnomAD high frequency variants, and ClinVar benign and pathogenic variants, 20 additional amino acid substitutions were generated by site directed mutagenesis of the AB598_yASS1 plasmid, and individual yeast strains harboring these variants were constructed by transformation with a PCR-amplified *pARG1-yASS1-tARG1-NatMX* cassette, as described above. These isolates, along with 12 standard curve variants from the original experiment chosen to cover a range of growth values, were phenotyped in a checkerboard format. The rest of the plate positions were filled in with the wild type *yASS1* strain, including edge positions, avoiding the necessity to normalize for neighbor growth and edge effects. Multiplicative normalization for plate effects was carried out and then a weighted least squares regression was used to estimate growth coefficients. The growth coefficients and standard errors were scaled to the original experimental dataset using the 12 standard curve variants via linear regression. The 20 variants measured in this second experiment are indicated in Table S4 as “Experimental Set 2”, while all other variants are indicated as “Experimental Set 1”.

## Results

### Establishing an *in vivo* ASS functional assay in yeast

We established a yeast-based assay to quantitatively measure the impact of amino acid substitutions on the activity of human ASS. This assay is based on the ability of human ASS to functionally replace (complement) its yeast ortholog (Arg1) in *S. cerevisiae*. We started with the human *ASS1* cDNA sequence (GenBank: NM_054012.4), which encodes the 412 amino acid ASS protein (*hASS1*). To maximize complementation, we also generated a codon-harmonized version of *ASS1* (*yASS1*) to more closely match the codon utilization frequency of yeast *ARG1*. A single copy of the human cDNA sequence or the yeast codon-optimized sequence was then integrated into a haploid yeast genome at the *ARG1* locus in place of the endogenous coding sequence and under the control of the endogenous *ARG1* promoter and terminator. This ensures that *hASS1* and *yASS1* are stably maintained as single copies and expressed at levels dictated by the native regulatory environment of the yeast ortholog of ASS1 (Figure 1A). Thus, in these strains, argininosuccinate synthetase activity is provided by human ASS in place of the native yeast enzyme. Since this enzymatic function is necessary for arginine biosynthesis, the activity of ASS is required for these yeast strains to proliferate in the absence of exogenous arginine. Therefore, growth on minimal medium lacking arginine is a quantitative measure of ASS enzyme function, allowing the impact of *ASS1* missense variants to be assessed at scale through a high-throughput growth assay.

**Figure 1.**
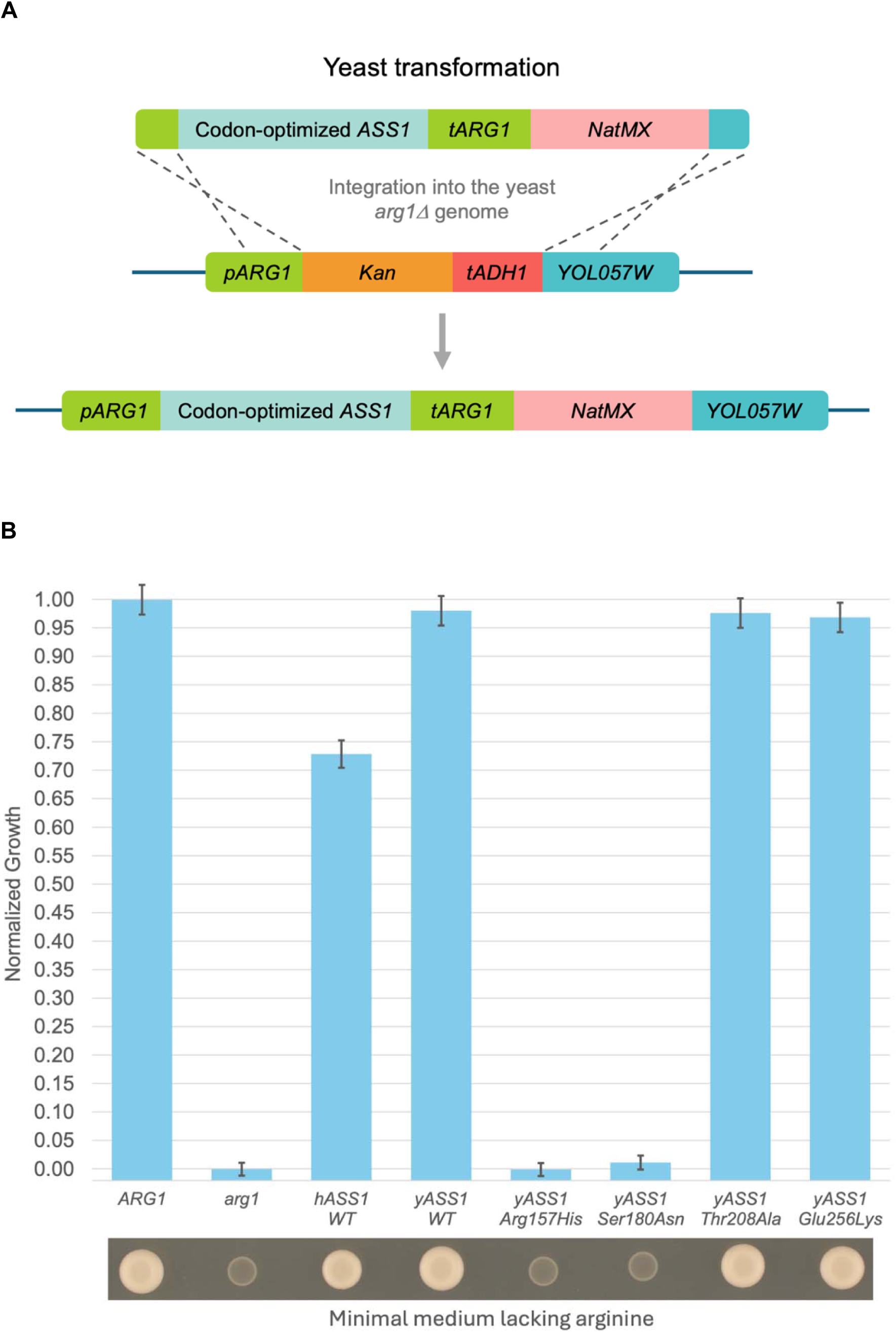
A yeast-based functional assay for human ASS amino acid substitutions. A) Yeast library construction. The *arg1Δ* strain was transformed with PCR products containing human *ASS1 (hASS1)* and either wild type or variant alleles of the yeast codon-optimized human *ASS1* coding sequence (*yASS1*), and a nourseothricin resistance drug marker (*NatMX*). Homology-directed integration at the *ARG1* locus places *hASS1* & *yASS1* under the control of the yeast *ARG1* promoter (*pARG1*) and terminator (*tARG1*). YOL057W represents the yeast gene downstream of the *ARG1* gene. B) Yeast growth in the absence of arginine as a quantitative assay for arginosuccinate synthetase function. Growth on minimal medium is calculated as the product of the area and the intensity from plates imaged after 3 days. Yeast strains harboring the native *ASS1* ortholog (yeast *ARG1*) grow robustly, but those harboring a precise gene deletion (*arg1*Δ) do not. Strains harboring wild type *hASS1*, wild type *yASS1* or benign variants (p.Thr208Ala or p.Glu256Lys) grow at 73%, 98%, 98% and 97% of *ARG1*, respectively, while strains harboring pathogenic variants (p.Arg157His or p.Ser180Asn) are unable to grow (0% & 1%, respectively). Growth estimates for each genotype +/- standard deviations are displayed relative to *ARG1* (set to 1) and *arg1Δ* (set to 0). Three independent isolates of each strain were assayed in triplicate.

To test the ability of hASS and yASS to functionally replace Arg1, we compared growth of the parental strain (FY4, expressing yeast *ARG1*) to that of the same strain background lacking any argininosuccinate synthetase activity (*arg1*Δ*0::NATMX*) or harboring the human *ASS1* expression constructs (*hASS1* or *yASS1*) at the *ARG1* locus as the sole source of enzyme activity. Each strain was grown under a non-selective condition (rich medium containing arginine), replica pinned to minimal medium lacking arginine, and grown at 30°C for 72 hours. As expected, FY4 grew robustly in the absence of arginine while the strain with the *arg1* deletion was unable to grow, displaying only the faint patch of cells deposited by the initial replica pinning (Figure 1B). Finally, human-based argininosuccinate synthetase (*ASS1*) was able to complement the *arg1* deletion; conferring 73% growth for human *ASS1* (*hASS1*), while yeast codon-optimized *ASS1* (*yASS1*) increased growth to 98% relative to the FY4 strain (Figure 1B). Since complementation was increased to 98%, all subsequent variant analysis was performed in the context of yeast codon-optimized *ASS1* (*yASS1*).

Our assay is designed to test the impact of amino acid substitutions on ASS activity. We expect that substitutions resulting from pathogenic missense variants will display reduced growth in our assay, while benign variants will result in little or no growth impairment. As a proof of principle, we tested the behavior of two known pathogenic and two known benign amino acid substitutions in our assay. The pathogenic substitutions correspond to the ClinVar pathogenic missense SNV NM_054012.4(*ASS1*):c.470G>A (p.Arg157His) and pathogenic/likely pathogenic missense SNV NM_054012.4(*ASS1*):c.539G>A (p.Ser180Asn). The common Arg157His sequence variant occurs in a highly conserved residue and leads to neonatal citrullinemia onset in homozygous patients [7]. *In vitro* studies have shown that the Arg157His substitution rendered ASS inactive, potentially through defects in aspartate recruitment into the substrate binding pocket [8]. Ser180Asn is another common sequence variant at a highly conserved residue that leads to classical citrullemia. *In vitro* studies have shown that it decreases the thermal stability of ASS and reduces affinity for both aspartate and citrulline [8]. When screened in our assay, the pathogenic Arg157His and Ser180Asn variants grew at 0% and 1% of wild type FY4 levels, respectively (Figure 1B). There are only four benign missense variants for *ASS1* in ClinVar and for our initial proof of principle test, we chose to assay the two variants with the highest population frequency reported in gnomAD, the SNVs NM_054012.4(*ASS1*):c.622A>G (p.Thr208Ala) and NM_054012.4(*ASS1*):c.766G>A (p.Glu256Lys), both of which affect surface residues. When screened in our assay, Thr208Ala and Glu256Lys grew at 98% and 97% of wild type FY4 levels, respectively (Figure 1B). Therefore, the activities of the tested ClinVar pathogenic and benign *ASS1* missense variants are consistent with expectations.

### Comprehensive analysis of the functional impact of SNV-accessible amino acid substitutions in ASS

Having validated our assay, we next applied it at scale to assess the functional impact of missense variants of *ASS1*. First, all unique amino acid substitutions that are accessible by a single nucleotide missense variant in the reference sequence of human *ASS1* were identified. In total, there are 2450 such amino acid substitutions. A library of *yASS1* derivatives, each encoding one of these amino acid substitutions, was then transformed into haploid yeast, replacing the native yeast *ARG1* ORF, as described above. To determine the identity of the codon change in each transformant, we sequenced the entire *yASS1* ORF using Oxford Nanopore sequencing. Transformants with secondary missense, indel, or nonsense mutations were removed from the final dataset.

We next assessed the growth of each individual isolate on selective solid medium plates (lacking arginine), at scale, using a gridded 8 x 12 array format, with robotic pinning from nonselective liquid medium. After 72 hours, plates were imaged and growth was measured as the sum of pixel intensities within each growth patch, as described in [4]. Normalization was then carried out to account for systematic plate-to-plate, edge, and neighbor effects on growth. The results were highly reproducible; for variants with more than one isolate, the correlation between the variant growth estimate (all isolates) and the individual isolate growth estimates had an r^2^ value of 0.947. A small number of variants (1.7%) were identified that displayed inconsistent growth between independent isolates, and these variants were removed from further analysis. Finally, a linear model was used to estimate the effect on growth of each tested amino acid substitution in ASS. The final growth values were then normalized so that growth of wild type *yASS1* was set to 1.0 and the *arg1* deletion strain was set to 0 (Table S4). The final number of unique amino acid substitutions functionally assessed was 2193, representing 90% of all SNV-accessible substitutions (Figure 2). A median of 5 isolates were generated for each amino acid substitution and 81% of substitutions were represented by more than one isolate.

**Figure 2.**
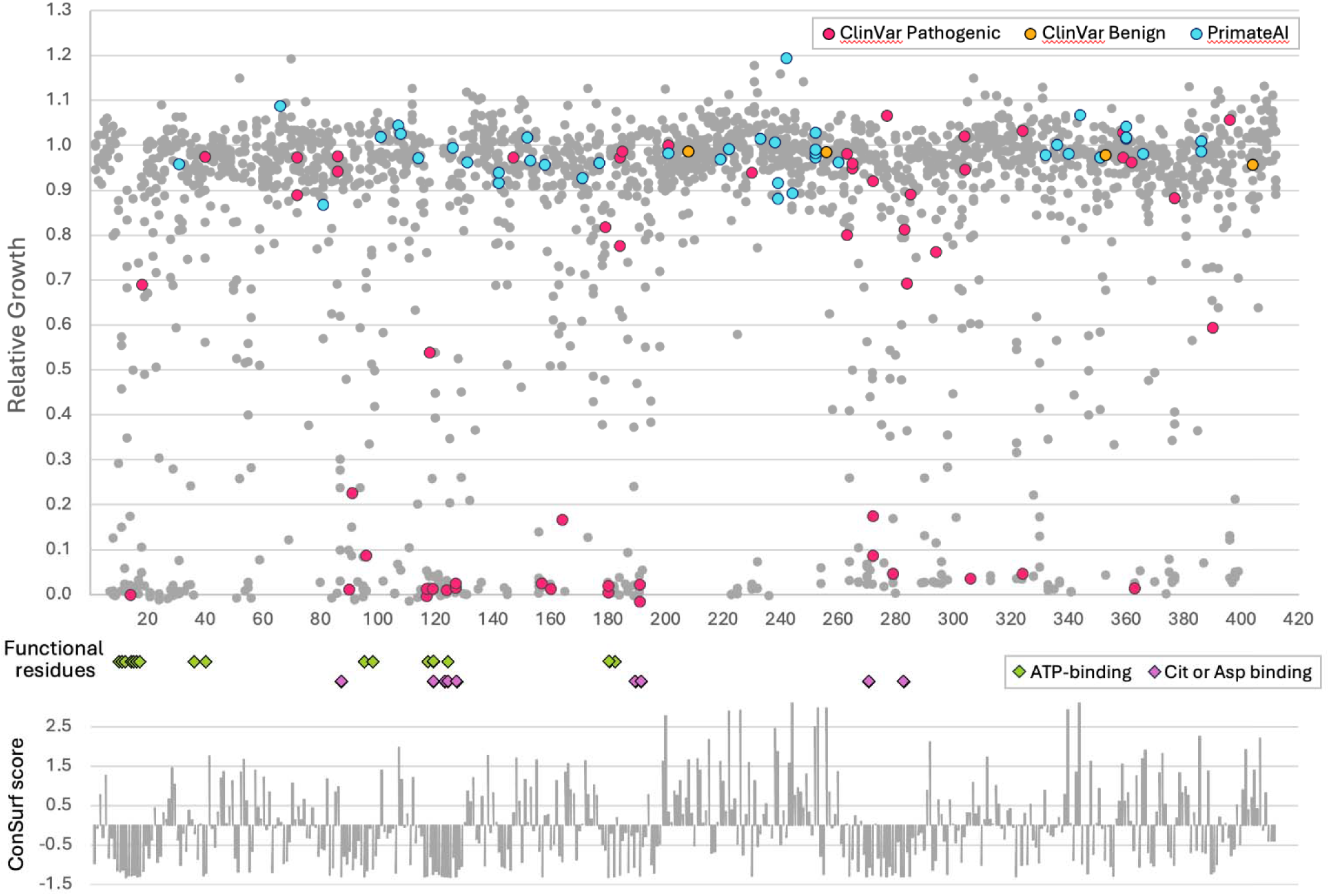
Relative growth of each amino acid substitution versus position in ASS, positions of ATP and substrate binding residues, and ConSurf conservation scores. Shown in magenta are amino acid substitutions corresponding to ClinVar pathogenic, pathogenic/likely pathogenic, and likely pathogenic variants. Shown in green are amino acid substitutions corresponding to ClinVar benign and likely benign variants. Shown in magenta are amino acid substitutions corresponding to ClinVar pathogenic, pathogenic/likely pathogenic, and likely pathogenic variants. Shown in green are amino acid substitutions corresponding to ClinVar benign and likely benign variants. Shown in light blue are amino acid substitutions corresponding to Primate.AI homozygous variants. Below are the positions of ASS1 functional residues (substrate binding), and ConSurf evolutionary conservation scores (PDB: 2nz2). More negative ConSurf scores indicate more conserved positions.

### Bimodal distribution of assay results identifies a large class of variants with severe loss of ASS function

The normalized growth scores of the 2193 variants tested in our functional assay display a bimodal distribution (Figure 3). The larger of the two peaks is centered around the wild type control (normalized growth=1.0), consistent with variants that display activity levels comparable to wild type in our assay. We classify these variants as “functionally unimpaired”. The smaller peak is centered around the null control (normalized growth = 0), consistent with variants exhibiting complete loss of function in our assay. We classify these variants as “functionally amorphic”. The remaining variants span a range of intermediate growth values, consistent with incomplete functional impairment, and are classified as “functionally hypomorphic.”

**Figure 3.**
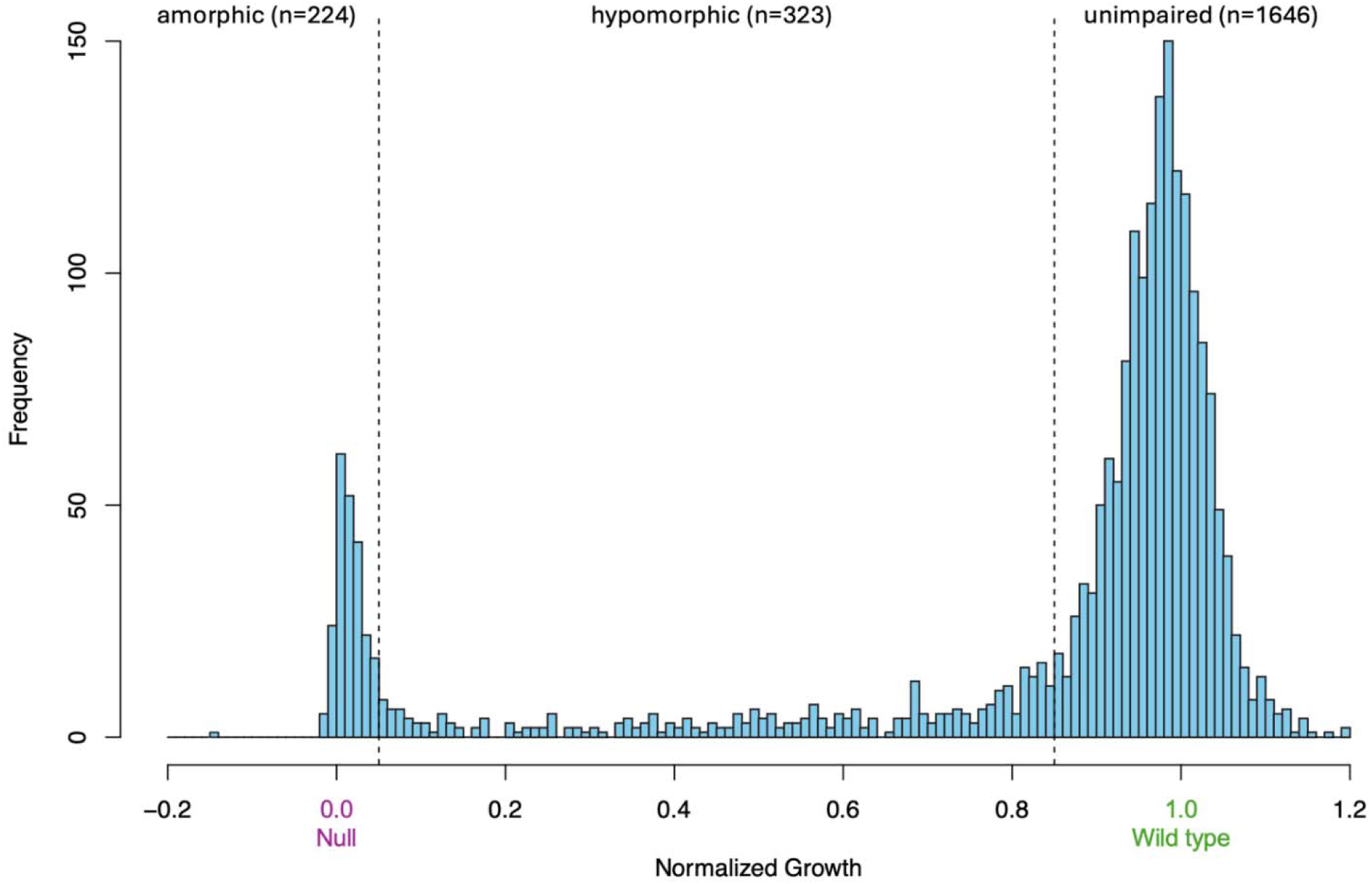
A bimodal distribution of normalized growth scores of the 2193 variants tested in our functional assay. Viewing variants ordered by normalized yeast growth and using thresholds set at 0.05 and 0.85, we classify 224 variants as functionally amorphic (<0.05), 323 variants as functionally hypomorphic (0.05-0.85), and 1646 variants as functionally unimpaired (>0.85).

We conservatively defined the lower boundary of the functionally unimpaired peak at 0.85 and the upper boundary of the amorphic peak at 0.05. Using these thresholds, the variants were classified as functionally amorphic (n= 224), hypomorphic (n= 323), and unimpaired (n= 1646) (Figure 3). The bimodal distribution of scores observed in our study, with peaks associated with wild type and null controls, is commonly seen in large-scale functional assays of protein function [9, 10].

### Assay results are consistent with evolutionary conservation and functionally important structural features

Human ASS is primarily expressed in the cytoplasm of hepatocytes where it functions as a homotetramer composed of a dimer of dimers (Figure 4A). Each 46 kDa monomer is made up of three domains: the first two, a nucleotide-binding domain and a synthetase domain, are positioned to form an active site pocket, while the third domain, a C-terminal helix involved in oligomerization, is separated from the other two domains by an extended loop (Figure 4B). ASS binding of ATP, citrulline, and aspartic acid occurs in the single active site pocket of each monomer. The dense set of amino acid substitutions characterized in our assay, with a median of 5 tested at each position in ASS, allowed us to compare our results with evolutionary conservation and structure-function relationships. We expected that substitutions at functionally critical residues would impair ASS activity.

**Figure 4.**
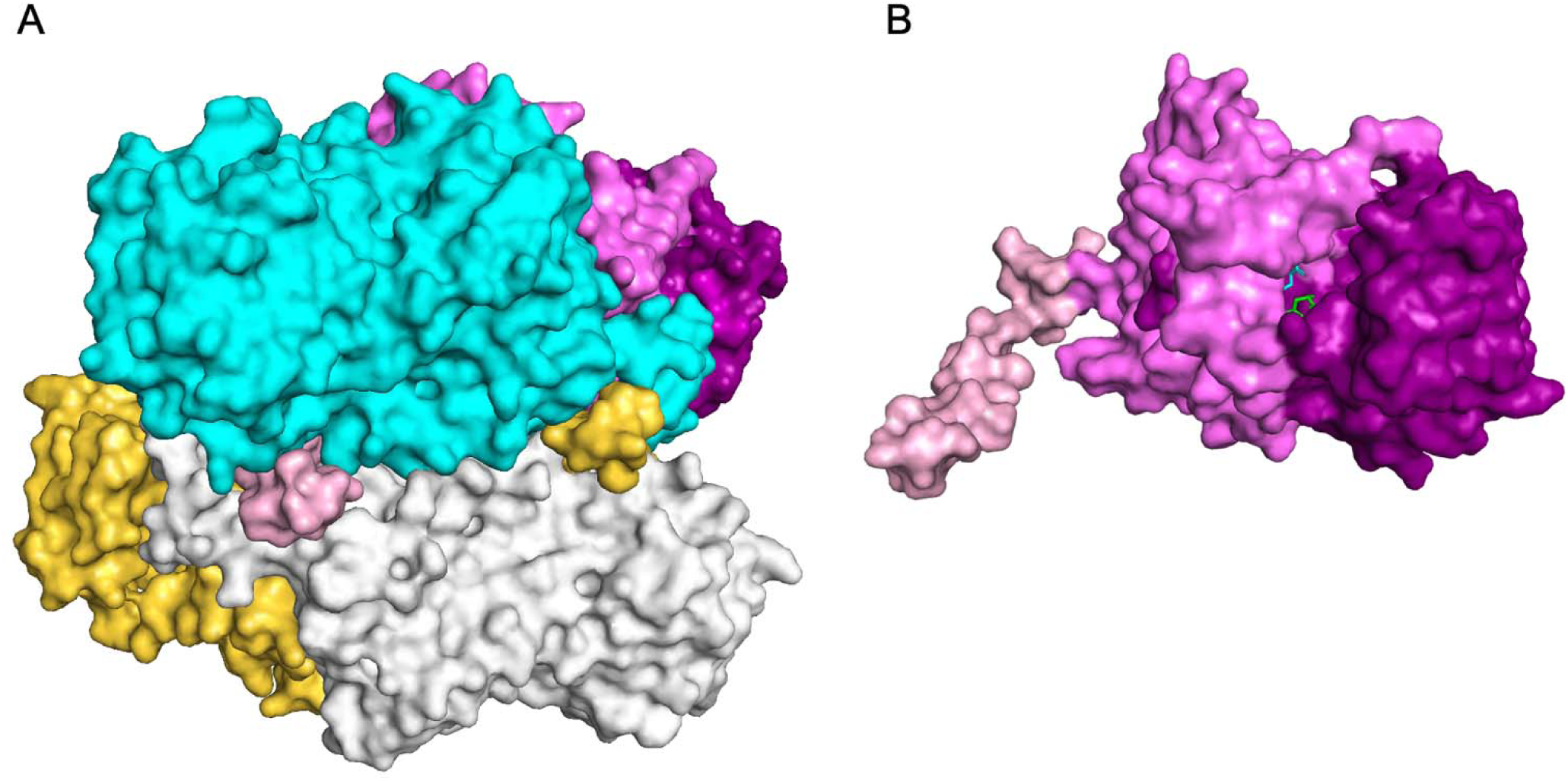
3-D Structure. A) The ASS homotetramer, B) Single monomer with its three domains displayed in different shades: nucleotide-binding domain (purple), synthetase domain (violet), and C-terminal helix (light pink). Substrates aspartate (green) and citrulline (cyan) are shown in the catalytic binding cavity.

ASS is an ancient enzyme, evolutionarily conserved from bacteria to mammals, and thus we expect amino acid substitutions that impair ASS function will be more likely to occur at highly conserved residues. To test this hypothesis, we compared variant growth in our functionally assay to evolutionary conservation, using the ConSurf database [11] to quantify the conservation of each residue in the human *ASS1* protein sequence (Figure 2). Consistent with our expectation, 73.2% (164/224) of amorphic substitutions (growth < 0.05 in our assay) occur at conserved residues (ConSurf relative conservation <0) which is significantly higher (Fisher’s Exact Test, p = 5.0 x 10^-11^) than the 50.1% (825/1646) of unimpaired substitutions (growth ≥ 0.85 in our assay) that occur at conserved residues (Figure 2).

Only one crystal structure of human ASS has been published to date, showing the enzyme bound to citrulline and aspartate, but not ATP [12]. Based on the authors’ analysis, the following residues come into direct contact with the substrates: citrulline is bound within the active site pocket through interactions with residues Tyr87, Asn123, Arg127, Ser189, Glu270, and Tyr282, while aspartate also interacts with Asn123, as well as Thr119. Additional analysis by Diez-Fernandez et al. [8] suggests that Glu191 is also involved in the binding of citrulline and that Asp124 participates in the binding of aspartate.

We predicted that amino acid substitutions made at these active site residues would be highly deleterious and our data are consistent with this expectation. A significantly higher proportion of variants at the nine cited active site residues display reduced activity in our assay (growth < 0.85), relative to all variants (90.7% vs 24.9%, chi-square goodness of fit test p = 5.4 x 10^-29^). In particular, the active site variants were highly enriched for severely impaired function in our assay (<0.05 growth) relative to all variants (61.1% vs 10.2%) (Figure 5A & Table S5). Looking in more detail at the behavior of individual active site residues, we find Thr119, Asn123, Asp124, Arg127, Glu191, and Glu270 to be particularly sensitive to amino acid substitutions. These results show strong agreement with enzymatic activity assays performed by Diez-Fernandez et al [8] where seven variants (Thr119Ile, Asp124Asn, Arg127Trp, Arg127Gln, Arg127Leu, Glu191Gln, and Glu270Gln) in five of these sensitive active site residues, which have been seen in patients with Citrullinemia type I, showed no activity in their assay.

**Figure 5.**
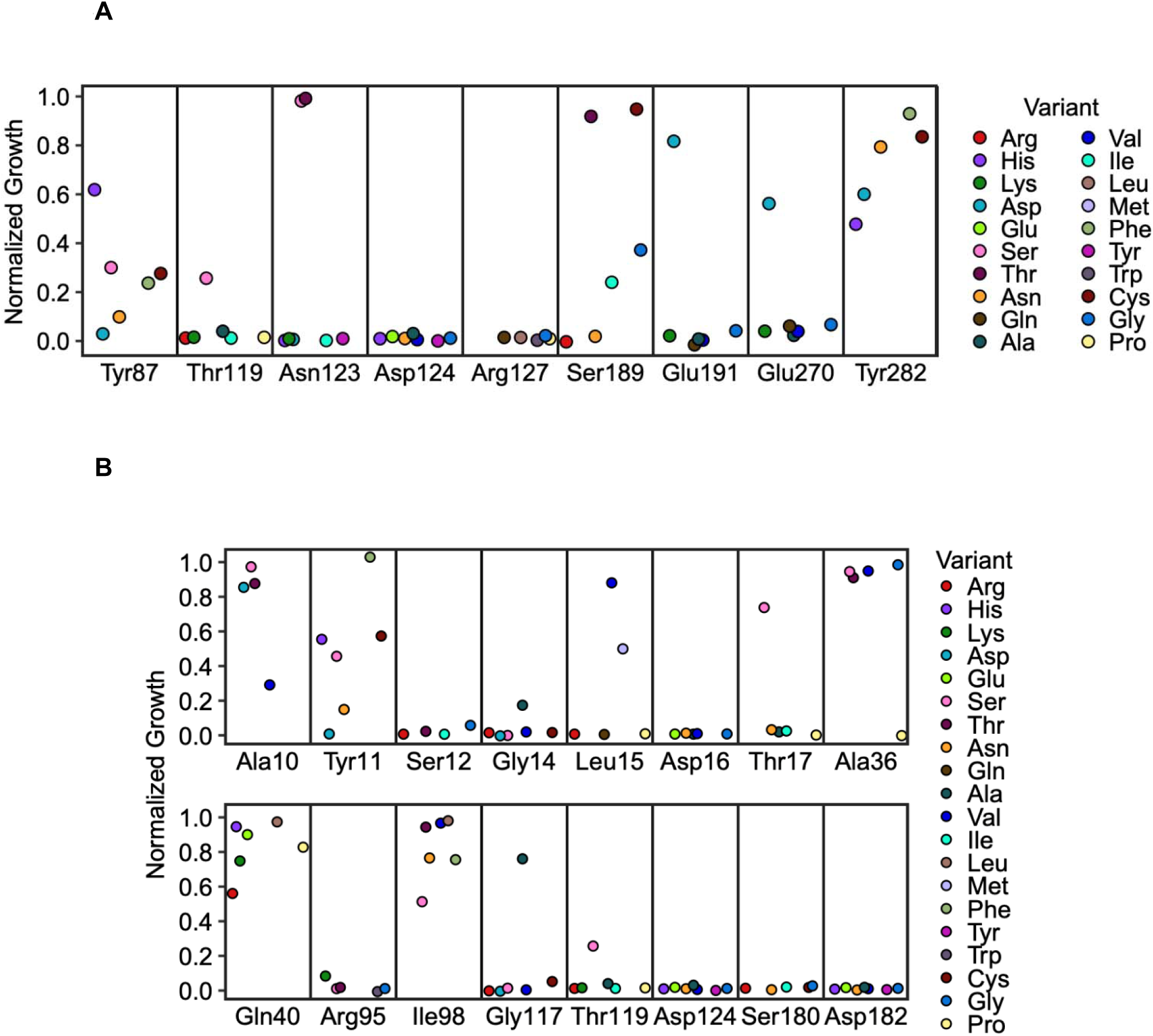
Effect of amino acid substitutions at functionally important residues. The distribution of variant growth scores for residues in ASS involved in A) citrulline and aspartate binding and B) ATP binding. Each circle is colored according to the amino acid that is substituted as shown (variant).

The residues involved in ATP binding in human ASS are not very well characterized. The single published crystal structure of human ASS, bound to both the citrulline and aspartate substrates, does not include ATP [12]. Even after extensive efforts, those authors reported that no satisfactory crystals were obtained with both substrates and ATP. Therefore, at this time, the identity of the residues of human ASS participating in the binding of ATP has been inferred based on crystal structures of argininosuccinate synthetase orthologs from *Escherichia coli* and *Thermus thermophilus*, which have been crystalized with bound ATP. Although orthologs display considerable primary sequence diversity, with *Homo sapiens* argininosuccinate synthetase sharing 26% and 52% sequence identity with *Escherichia coli* and *Thermus thermophilus*, respectively, the overall structure of the tetrameric enzyme is highly conserved.

From studies of *E. coli* argininosuccinate synthetase in complex with ATP, and with ATP plus citrulline, Lemke et al. [13] identified residues that contact ATP corresponding to human ASS residues Ala10, Ser12, Leu15, Asp16, Thr17, Ala36, Gly117, Thr119, Asp124, and Asp182. Concurrently, crystal studies of argininosuccinate synthetase from *T. thermophilus* [14, 15] identified ATP-binding residues corresponding to human ASS residues Ala10, Ala36, Arg95, Ile98, Gly117, and Ser180, as well as Tyr11, Ser12, Gly14, Leu15, and Asp16 of the P-loop (residues 11-18), which is an ATP binding domain found in the family of “N-type” ATP pyrophosphatases that includes argininosuccinate synthetase [16]. Lastly, Karlberg et al proposed that Gln40 forms a hydrogen bond to the adenine group of ATP [12].

We predicted that residues directly involved in ATP binding would be highly sensitive to amino acid substitutions. 83.0% of variants at the 16 inferred ATP binding residues display reduced activity in our assay (growth < 0.85), compared to 24.9% of all variants (chi-square goodness of fit test p = 2.8 x 10^-36^) and 61.4% of variants affecting ATP-binding residues are amorphic versus 10.2% of all variants. We found Ser12, Gly14, Asp16, and Thr17 within the P-loop, as well as Arg95, Gly117, Thr119, Asp124, Ser180, and Asp182 to be particularly sensitive to amino acid substitutions, consistent with their inferred role in ATP binding. In contrast, Ala10, Ala36, Gln40 and Ile98 were more tolerant to substitution (Figure 5B & Table S5). These results are also in agreement with enzymatic activity assays performed by Diez-Fernandez et al [8] where six variants (Arg95Ser, Gly117Ser, Thr119Ile, Asp124Asn, Ser180Asn, and Ser180Ile) found in five of these sensitive ATP-binding residues showed little to no activity in their assay.

To identify additional sensitive residues, we mapped all positions where ≥50% of tested variants were amorphic (Table S6). Out of a total of 29 such sensitive residues, 22 (76%) map to the edge or within the active site pocket, with 15 of those residues participating in direct interactions with ATP, citrulline, or aspartate, as described above, while the remaining residues (Ala118, Lys121, Gly122, Gly156, Arg157, Leu160, and Met377) may also be important for catalytic function, perhaps by scaffolding the structure of the active site pocket and the conformation of residues that directly interact with ligands. Of the 7 other sensitive residues (24%) which do not map to the active site pocket, the majority (Phe333, Cys337, Arg363, and Arg398) are likely to participate in oligomerization. Phe333 and Cys337 of one dimer, may interact with the Cys337 and Phe333 of the other dimer, respectively, via a hydrophobic interaction between the phenylalanine aromatic ring and the sulfur-containing cysteine chain via aromatic-thiol hydrogen bonding, to promote oligomerization of the homotetramer. Arg363 is likely involved in folding with an ionic interaction with D296 on the same monomer [12], as well as in dimer formation via a salt bridge with Asp267 of the other monomer. Arg398 in the C-terminal helix is likely involved in dimer formation in a charged interaction via a salt bridge with Glu330 of the other monomer, and these two residues are also in close proximity to Arg304 and Glu305 in the neighboring dimer, which may also contribute to oligomerization of the homotetramer. The remaining 3 residues (Gly230, Arg279, and Gly324) are buried and may play structural roles in the proper folding of each monomer.

In summary, analysis of variant effects in a structural context demonstrates that our assay successfully identifies residues critical to catalytic function, substrate binding, and oligomerization. Our assay results are consistent with the expectation that amino acid substitutions are detrimental to function in residues critical for catalysis. In addition, our data suggest that variant effect maps can reveal functionally important positions that are not obvious from structural models alone.

### Functional assay provides PS3_supporting level evidence towards pathogenicity for *ASS1* variants

To assess the potential of our assay to provide supporting evidence for variant pathogenicity or benignity according to the American College of Medical Genetics and Association for Molecular Pathology (ACMG/AMP) guidelines, we first examined variants that are associated with pathogenic or benign clinical annotations in the ClinVar database [17]. At the time of writing, there were 59 *ASS1* missense variants with pathogenic or likely pathogenic annotations and 4 missense variants with benign or likely benign annotations. We tested the impact of the amino acid substitutions corresponding to 56 pathogenic variants and all four benign variants (Figure 6).

**Figure 6.**
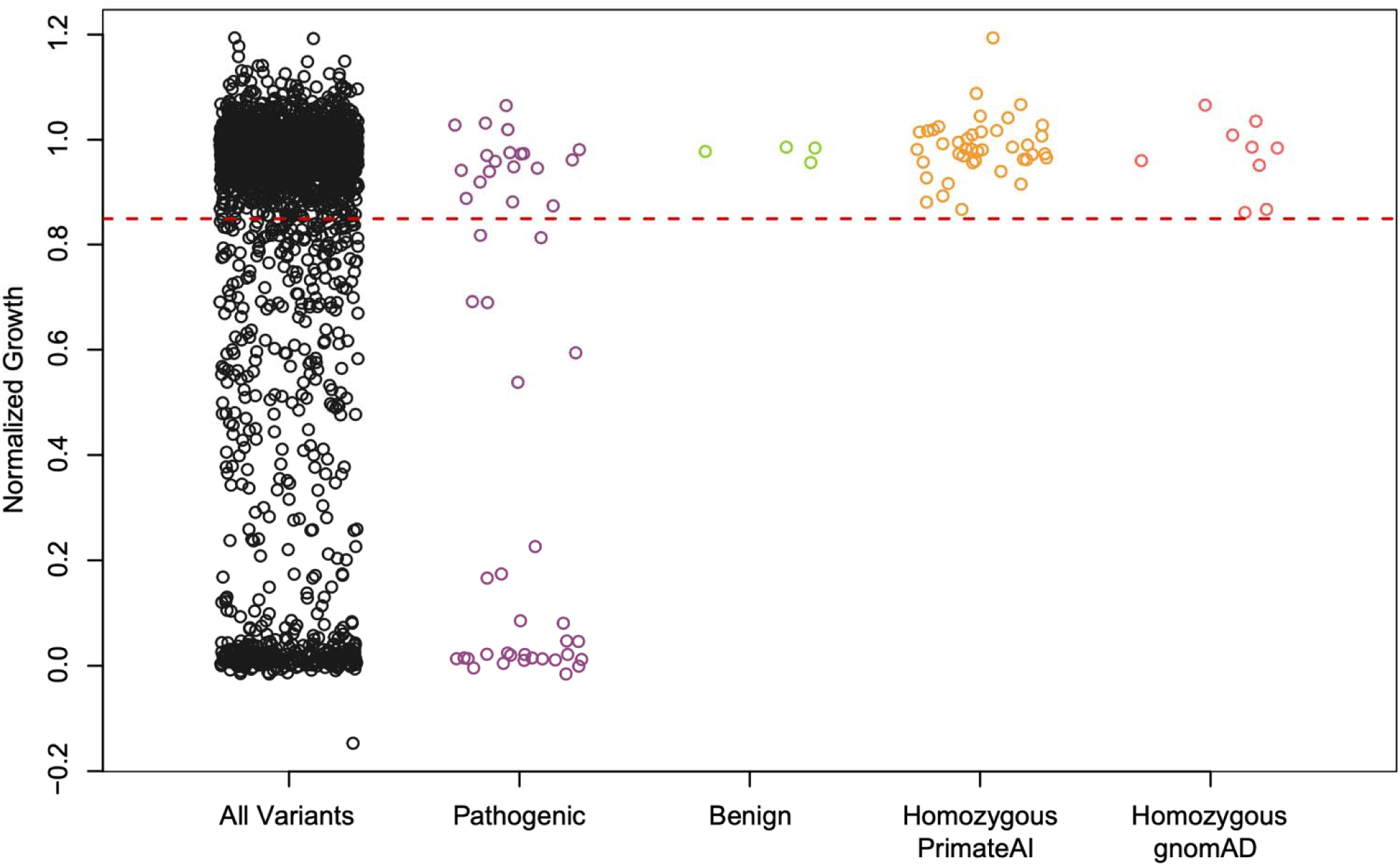
Strip charts of normalized growth for all amino acid substitutions and those corresponding to ClinVar, homozygous Primate.AI, and homozygous gnomAD classification groups

In the range of our assay associated with reduced or null variant activity (normalized growth <85%), substitutions corresponding to 34 pathogenic and 0 benign variants were observed. This is consistent with reductions in ASS function corresponding to <85% growth in our assay being associated with disease in humans. The associated Odds of Pathogenicity (OddsPath) for this range of the assay (2.43) is sufficient to provide “supporting” strength evidence under the ACMG/AMP PS3 criterion [18] for the 547 amino acid substitutions with <85% normalized growth in our assay. This includes 25 substitutions that correspond to ClinVar missense variants currently classified as variants of uncertain significance (VUS).

A major limitation in functional variant interpretation for many genes associated with rare genetic disorders, including *ASS1*, is the small number of ClinVar missense variants with benign annotations. For instance, *ASS1* has only four ClinVar benign missense variants, all of which displayed growth >95% in our assay. A possible solution to the rarity of ClinVar benign variants is to use data from non-human primates. Because of the short evolutionary distances between humans and other primates, homozygous variants observed in other primates are highly likely to be benign in humans. The PrimateAI-3D database (previously PrimAD) [19] has collated variant information from 809 individuals of 233 non-human primate species and demonstrated that variants observed in these individuals are ∼100 fold more likely to be classified with high confidence as benign/likely-benign in ClinVar than pathogenic/likely-pathogenic [20].

On this basis, we examined *ASS1* amino acid substitutions tested in our assay that correspond to homozygous missense variants seen in at least one individual non-human primate in PrimateAI-3D. There were 41 such variants, all of which exhibited growth above our 85% cutoff, consistent with reduced fitness (disease association) of variants falling below this threshold, and purifying selection, in primates. If we treat PrimateAI variants as benign (solely for OddsPath calculation purposes), the OddsPath for growth <85% rises to 27.32, reaching the full PS3 level of evidence for pathogenicity under current guidelines [18]. Interestingly, among all nine of the human *ASS1* missense variants observed as homozygotes in gnomAD which were tested as amino acid substitutions in our assay, the lowest observed growth value was 0.86 (Ala258Val) consistent with the lower boundary of the PrimateAI variants which was a growth value of 0.87 (Ala81Thr) (Figure 6).

While growth below 85% in our assay showed strong enrichment for pathogenic vs benign variants, 22 of the 56 pathogenic missense variants still displayed growth above this threshold. To investigate this, we examined the spatial positioning of the pathogenic variants that we tested. The majority of the 56 variants fall into three main classes; those located within the active site pocket (23 variants in 16 residues) (Figure 7A), those falling on a large monomer-monomer interface (11 variants in seven residues) (Figure 7B), and those likely impacting protein stability and folding (20 variants in 16 residues) (Figure 7C). The sensitivity of our assay varied strongly between these classes with 20/23 active site variants, 10/20 stability variants and only 2/11 interface variants displaying reduced growth (<85%). Interestingly, the two interface variants that did show reduced growth occur at a residue (Arg363) that also has a role in protein folding (Table S7). Therefore, our assay appears to be very sensitive to variants affecting active site function, variably sensitive to variants impacting folding, and insensitive to variants located on the dimerization interface.

**Figure 7.**
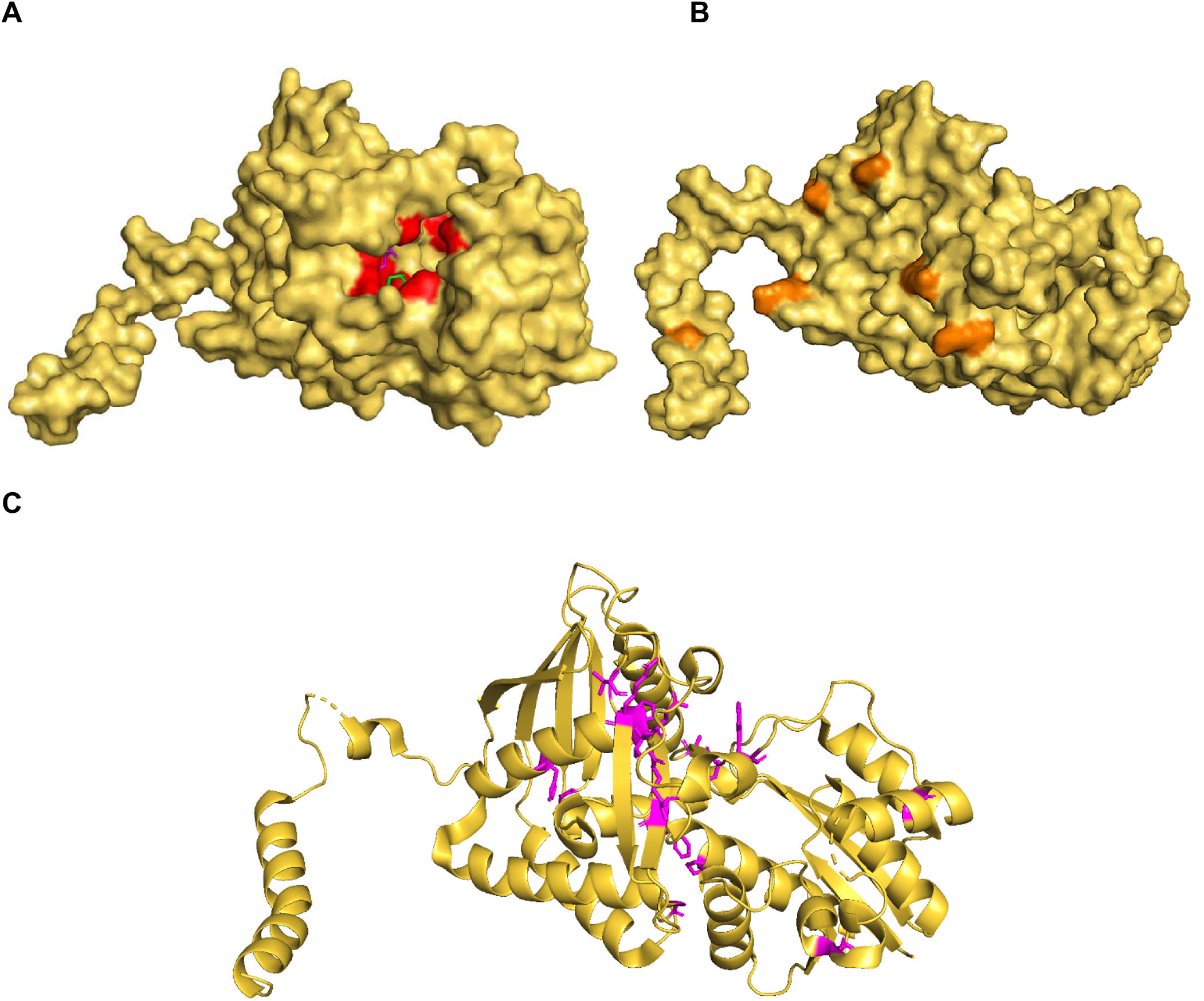
ClinVar pathogenic active site variants & pathogenic dimerization variants. Single monomer in yellow highlighting A) residues corresponding to ClinVar pathogenic variants involved in active site formation/function in red, B) residues corresponding to ClinVar pathogenic variants at the dimerization interface in orange, and C) residues corresponding to ClinVar structural pathogenic variants in magenta.

These results suggest that active sites residues are necessary for ASS activity in both human and yeast cells, but that dimerization residues and some folding residues may be more important for protein function in the environment of human cells. Alternatively, high-growth pathogenic variants may cause only mild enzymatic loss-of-function, and the resulting residual activity, combined with yeast-specific expression levels, may be sufficient for normal growth in our assay. Supporting this interpretation, many high-growth pathogenic variants are associated with late-onset or milder clinical presentations [7, 21–31]. Under this model, the reason that we successfully detect pathogenic active site variants, but not dimerization variants, is that active site variants generally have a larger effect on protein function.

Based on these findings and given that 22/56 pathogenic variants show ≥85% growth, we conclude that growth above this threshold should not be used as evidence toward benignity. Conversely, the variants with <85% growth are likely to cause a strong loss of ASS activity and are strongly associated with pathogenicity.

### Regions from two subunits contribute to each ASS active site which allows intragenic complementation to occur

CTLN1 is an autosomal recessive disorder. As such, disease manifestation depends on the combined activity of both *ASS1* alleles. If this combined activity falls below a critical threshold, CTLN1 is expected to occur.

In a previous study of human *PSAT1*, we demonstrated that pairwise allele activity could be accurately predicted from single variant estimates using an additive model [32]. This represents a powerful approach to leveraging large scale functional single variant datasets, such as the one presented here, for genotype-based disease risk predictions. However, this additive framework fails in the presence of epistasis between alleles, including dominant negative effects or intragenic complementation.

Intragenic complementation occurs when two deleterious alleles, each displaying severe loss of function on their own, together restore significant protein activity in a compound heterozygous state. In 1964, Crick and Orgel [33] proposed a model to explain this phenomenon in homomultimeric enzymes, where protein function can be restored when non-functional variants are sequestered into a subset of the active sites, leaving the remaining active sites functional (Figure 8A). This variant sequestration model requires the enzyme is a homomultimer with multiple active sites and that each active site is composed of residues contributed by multiple monomers.

**Figure 8.**
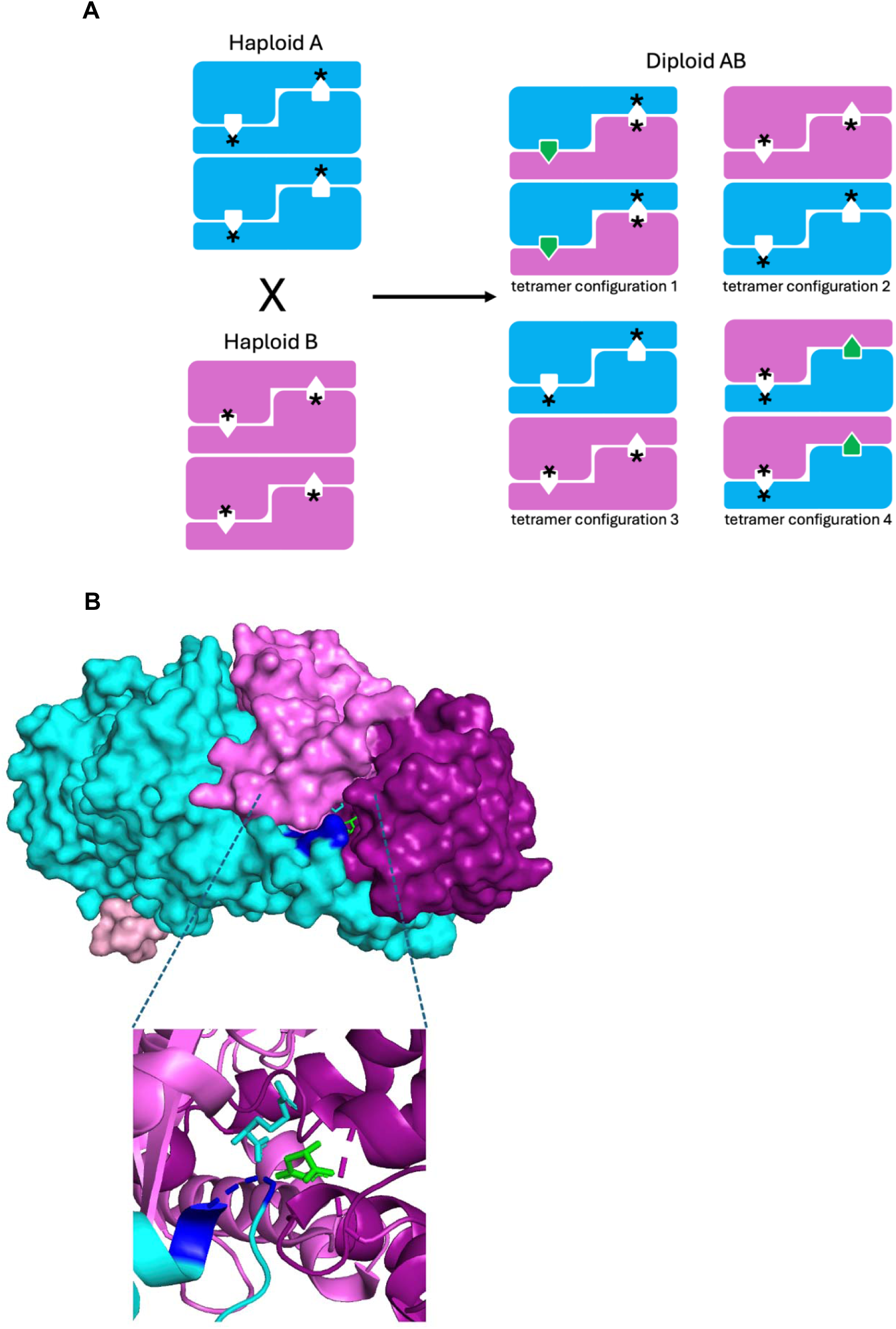
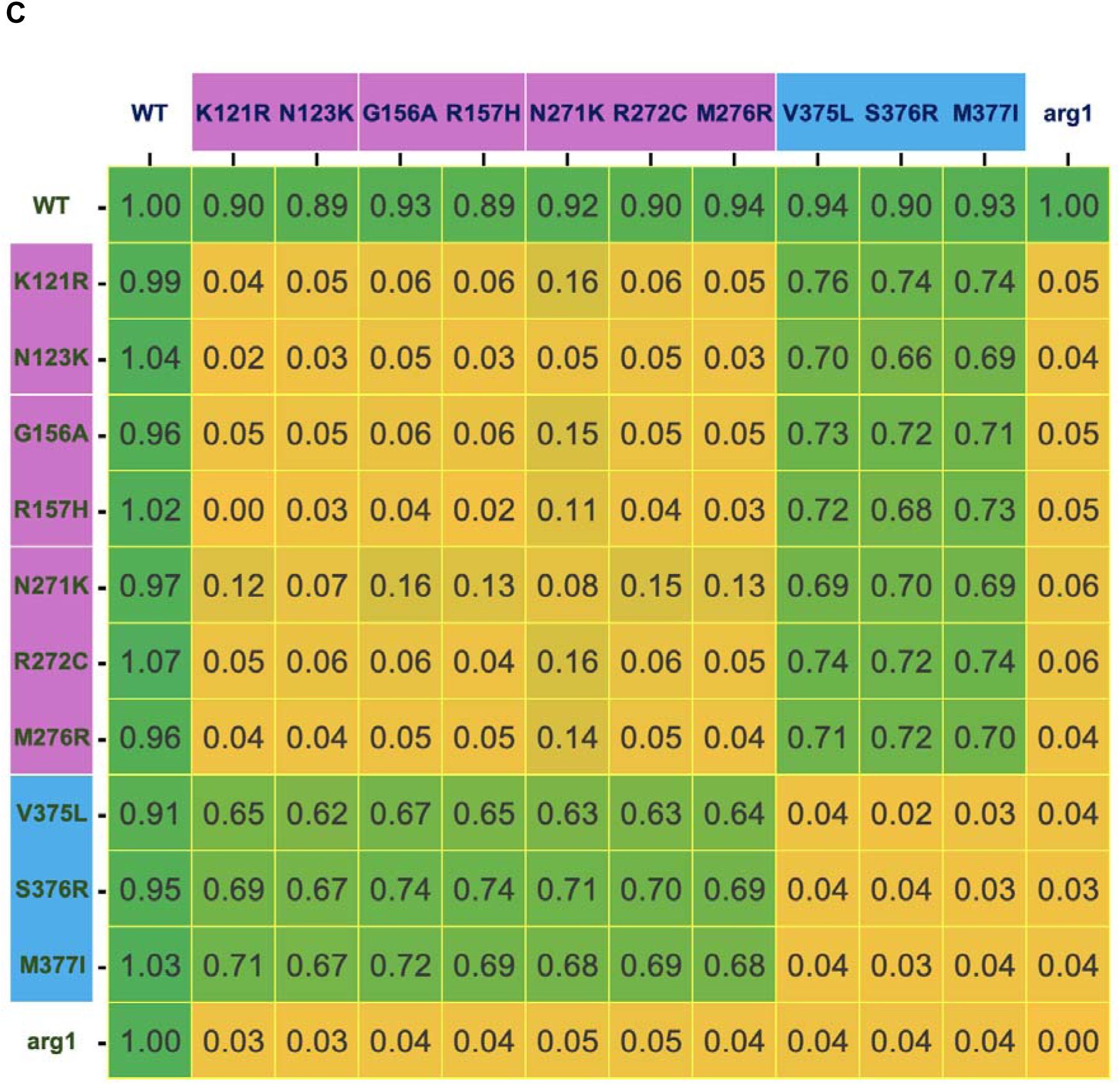
Intragenic complementation. A) Intragenic complementation model with graphical representations of two separate ASS amorphic active site variants tetramerizing to form both active and inactive catalytic sites. B) 3-D dimer structure illustrating two monomers contributing to a compound active site with an enhanced view. Residues 375-378 are shown in blue, and substrates aspartate & citrulline are shown in green and cyan, respectively. C) Growth scores of an intragenic complementation assay with normalized scores of diploid growth of all combinations tested between amorphic variants at two amino acid positions. Active site variants are labeled in purple boxes, extended arm variants are labeled in blue boxes.

ASS is a homotetramer with four active sites, suggesting potential for complementation, but to our knowledge, evidence of inter-subunit contribution to individual active sites has not previously been reported. However, structural analysis identified a region of the extended loop separating the synthetase domain from the C-terminal helix that comes into close proximity with the active site pocket of an adjacent monomer. Specifically, residues 375-378 in this loop lie adjacent to residues 155-157 at the mouth of the active site pocket (Figure 8B). Residues Gly156 and Arg157 are highly sensitive to amino acid substitutions, as is Met377, suggesting that these may form part of a compound active site spanning two monomers (active site pocket and extended loop). Interestingly, both Met377 and Gly156 lie in disordered regions in the human ASS crystal structure [12].

To test this model, we hypothesized that amorphic (severe loss-of-function) variants in the extended loop and the active site pocket would complement each other *in trans*, as per Crick and Orgel’s variant sequestration model. Specifically, we tested amorphic loop region variants Val375Leu, Ser376Arg, and Met377Ile against amorphic active site variants Lys121Arg, Asn123Lys, Gly156Ala, Arg157His, Asn271Lys, Arg272Cys, and Met276Arg. Diploid yeast strains were constructed by mating haploids of opposite mating type, each harboring one variant. We observed that all homozygous combinations of these amorphic variants showed persistent loss of function, as did all combinations involving two variants from the same region (either both loop or both active site). However, compound heterozygotes with one variant in the active site and one in the extended loop consistently restored substantial ASS activity (Figure 8C). These results provide the first direct evidence of intragenic complementation in *ASS1*, and identify residues Val375, Ser376, and Met377 in the extended loop as critical contributors to a shared or compound active site along with residues in the canonical active site pocket.

## Discussion

Citrullinemia type I (CTLN1) is a rare but potentially life-threatening metabolic disorder. As a monogenic disease caused by pathogenic variants in the *ASS1* gene, CTLN1 is amenable to detection through genetic sequencing. Newborn sequencing of *ASS1* has the potential to reliably identify at-risk individuals presymptomatically. This genetic information is highly clinically actionable. In individuals with appreciable ASS deficiency, early treatment with nitrogen-scavenging agents, arginine supplementation, and strict dietary management can prevent catastrophic outcomes, including irreversible brain injury and death. Early intervention is further associated with improved developmental outcomes. For these individuals, sequencing offers key advantages over biochemical testing, providing a more definitive diagnosis, and opening a critical window for clinical intervention before a hyperammonemic crisis.

In the near future, early diagnosis of individuals with severe ASS deficiency may also enable timely access to gene therapies, which hold the potential to provide durable or even curative treatment for CTLN1. This field is advancing rapidly, as illustrated by recent reports of successful interventions for other urea cycle disorders. In the OTC-HOPE trial [34] gene therapy for ornithine transcarbamylase (OTC) deficiency led to discontinuation of ammonia-scavenging drugs and transition to age-appropriate protein intake, with sustained normal ammonia levels and no hyperammonemic crises through six months of follow-up. Similarly, a customized CRISPR-based base-editing therapy for a 6-month-old boy with CPS1 deficiency resulted in improved protein tolerance and a substantially reduced need for ammonia scavengers [35]. These cases underscore the growing clinical feasibility of molecular cures for inborn errors of metabolism.

Newborn sequencing also plays a critical role in identifying individuals with partial, rather than severe, ASS impairment, who can be missed by standard newborn biochemical screening due to near-normal metabolite levels. These individuals remain at risk for late-onset disease, which has the potential to be lethal [36], but often presents with vague or non-specific symptoms such as psychiatric disturbances [35] that may not be recognized as manifestations of ASS deficiency. Biochemical testing alone may fail to detect their disease, delaying recognition and appropriate care. In contrast, sequencing has the potential to unambiguously identify these at-risk individuals early in life, enabling their condition to be effectively managed through lifelong dietary control and close monitoring of environmental triggers. In the future, gene therapy may also become appropriate for individuals with partial deficiency, if the benefit-risk profile proves superior to that of existing management strategies for this population.

Projects such as the GUARDIAN study [37] and the Generation Study (Genomics England) [38] are evaluating the effectiveness of DNA sequencing for newborn screening of genetic disorders. Within these programs, sequencing of *ASS1* has been prioritized because accurate and early identification of both severe and partial forms of ASS deficiency enables effective clinical intervention. However, in common with other genes, interpretation of missense variants in *ASS1* remains a major roadblock in sequencing-based diagnosis. As of writing, ClinVar includes 59 pathogenic missense variants, 4 benign variants, 33 variants with conflicting calls, and 117 classified as variants of uncertain significance (VUS) which are not clinically actionable. With the continued expansion of genome sequencing, particularly if sequencing based newborn screening becomes widespread, the number of VUS in *ASS1* is likely to increase.

Functional assays have the potential to resolve VUS by providing direct evidence of biological impact, a critical capability for rare disorders like CTLN1, where case-based data are often sparse. We therefore developed a large-scale yeast-based functional assay for ASS activity and quantified the functional impact of 2,193 amino acid substitutions, representing 90% of all single-nucleotide variant (SNV)-accessible substitutions. Of the 2,193 substitutions, 224 were functionally amorphic (<5% activity), and an additional 323 were hypomorphic (5%–85% activity). When applied within the ACMG/AMP OddsPath framework [18], and based on the behavior of ClinVar benign and pathogenic variants, growth below 85% in our assay meets the threshold for PS3_supporting evidence of pathogenicity. Therefore, for the 547 amino acid substitutions that fell below this functional threshold, our assay provides evidence of pathogenicity at this level for the corresponding human missense variants.

A major limitation of relying on ClinVar for assay calibration is the small number of annotated benign missense variants typically available, which constrains the maximum evidence strength that functional data can support. In our case, the presence of only four benign or likely benign ClinVar variants prevented our functional evidence from exceeding the “Supporting” level. To address this, we incorporated 41 SNV-accessible amino acid substitutions corresponding to missense variants observed as homozygotes in other primates. All 41 were functionally unimpaired in our assay. These variants, segregating at appreciable frequency in other species with no disease phenotype, serve as an orthogonal calibration set for benign missense variation in *ASS1*. When used (solely) for assay calibration, they increase the functional evidence strength for pathogenicity from PS3_supporting to full PS3, substantially expanding the clinical utility of our dataset for variant interpretation. Integration of this high-resolution functional evidence into diagnostic pipelines, alongside population, segregation, and computational data, has the potential to support reclassification of current *ASS1* VUS and facilitate earlier, more confident diagnoses of CTLN1.

While variant-level functional data calibrated with orthogonal benign controls enhances our ability to classify individual missense changes, clinical interpretation ultimately depends on genotype-level context, that is, the combined functional effect of both alleles present in a person. For autosomal recessive conditions like CTLN1, where two variants must act *in trans* to cause disease, pathogenicity is determined by their joint impact on protein function. If the combined activity of the two alleles falls below a critical threshold, disease manifests; as residual activity declines further, earlier onset and/or more severe forms of disease are expected.

The relationship between single-variant effects and their combined impact *in trans* is often assumed (explicitly or implicitly) to be additive. For example, two severe loss-of-function variants in compound heterozygosity are typically predicted to result in near-complete loss of function and correspondingly severe disease. However, in this study, we demonstrate that certain combinations of severe *ASS1* variants, when affecting distinct structural regions such as the active site and the extended inter-subunit loop, can restore substantial enzymatic activity in compound heterozygotes. This phenomenon, known as intragenic complementation, is a form of epistasis that results in combined allele activity higher than expected under an additive model.

Our data are consistent with Crick and Orgel’s classic model of intragenic complementation via variant sequestration in homomultimeric enzymes [33] and provide the first direct evidence that ASS active sites span multiple subunits. While single-variant functional scores are critical for classification, this finding illustrates how non-additive interactions between alleles can alter clinical interpretation. Intragenic complementation could obscure variant pathogenicity in compound heterozygotes, or conversely, partially rescue function in allelic combinations previously assumed to be deleterious. These findings highlight the need to consider allele pair context in clinical genetics, particularly for autosomal recessive conditions like CTLN1.

## Conclusions

Citrullinemia type I (CTLN1) is a rare but treatable disorder for which sequencing-based newborn screening holds significant potential. Early molecular diagnosis could enable timely intervention, improved developmental outcomes, and, in the future, access to curative gene therapies. However, the widespread adoption of genomic screening is likely to increase the number of identified ASS1 variants of uncertain significance, creating a major barrier to clinical actionability and timely diagnosis. To address this, we developed a high-throughput assay that provides functional evidence for the impact of 2,193 ASS1 missense substitutions. Based on ClinVar annotations alone, these data support PS3_supporting-level evidence for pathogenicity under ACMG/AMP criteria. However, when calibrated using orthologous benign variants observed as homozygotes in other primates, these data reach full PS3 strength, substantially enhancing their clinical utility. Additionally, we show that intragenic complementation between certain ASS1 loss-of-function variants can restore enzymatic activity, underscoring the importance of allele-pair context in clinical interpretation. These findings support the integration of functional and combinatorial data into diagnostic workflows for CTLN1.

## Acknowledgements

This work utilizes a small equipment grant from the National Urea Cycle Disorders Foundation.

## References

1. Ah Mew N, Simpson KL, Gropman AL, Lanpher BC, Chapman KA, Summar ML: Urea Cycle Disorders Overview. In: GeneReviews((R)). Edited by Adam MP, Feldman J, Mirzaa GM, Pagon RA, Wallace SE, Amemiya A. Seattle (WA); 1993.

2. Richards S, Aziz N, Bale S, Bick D, Das S, Gastier-Foster J, Grody WW, Hegde M, Lyon E, Spector E et al: Standards and guidelines for the interpretation of sequence variants: a joint consensus recommendation of the American College of Medical Genetics and Genomics and the Association for Molecular Pathology. Genet Med 2015, 17(5):405–424.

3. Rose M, Winston F, Hieter P: Methods in Yeast Genetics: A Laboratory Course Manual. Cold Spring Harbor, NY: Cold Spring Harbor Laboratory Press; 1990.

4. Lo RS, Cromie GA, Tang M, Teng K, Owens K, Sirr A, Kutz JN, Morizono H, Caldovic L, Ah Mew N et al: The functional impact of 1,570 individual amino acid substitutions in human OTC. Am J Hum Genet 2023, 110(5):863–879.

5. SRA – NCBI. [https://www.ncbi.nlm.nih.gov/sra/.] Sequence Read Archive data under SRA accession PRJEB91359. Accessed 3 July 2025.

6. GitHub [https://github.com/lacyk3/PyPl8.] PyPl8 Image analysis package for segmenting images of microbial communities and extracting quantitative features automatically. Accessed 7 July 2025.

7. Diez-Fernandez C, Rufenacht V, Haberle J: Mutations in the Human Argininosuccinate Synthetase (ASS1) Gene, Impact on Patients, Common Changes, and Structural Considerations. Hum Mutat 2017, 38(5):471–484.

8. Diez-Fernandez C, Wellauer O, Gemperle C, Rufenacht V, Fingerhut R, Haberle J: Kinetic mutations in argininosuccinate synthetase deficiency: characterisation and in vitro correction by substrate supplementation. J Med Genet 2016, 53(10):710–719.

9. Gronbaek-Thygesen M, Voutsinos V, Johansson KE, Schulze TK, Cagiada M, Pedersen L, Clausen L, Nariya S, Powell RL, Stein A et al: Deep mutational scanning reveals a correlation between degradation and toxicity of thousands of aspartoacylase variants. Nat Commun 2024, 15(1):4026.

10. Choudhury A, Fenster JA, Fankhauser RG, Kaar JL, Tenaillon O, Gill RT: CRISPR/Cas9 recombineering-mediated deep mutational scanning of essential genes in Escherichia coli. Mol Syst Biol 2020, 16(3):e9265.

11. The ConSurf Database [https://consurfdb.tau.ac.il/.] Accessed 7 July 2025.

12. Karlberg T, Collins R, van den Berg S, Flores A, Hammarstrom M, Hogbom M, Holmberg Schiavone L, Uppenberg J: Structure of human argininosuccinate synthetase. Acta Crystallogr D Biol Crystallogr 2008, 64(Pt 3):279–286.

13. Lemke CT, Howell PL: Substrate induced conformational changes in argininosuccinate synthetase. J Biol Chem 2002, 277(15):13074–13081.

14. Goto M, Nakajima Y, Hirotsu K: Crystal structure of argininosuccinate synthetase from Thermus thermophilus HB8. Structural basis for the catalytic action. J Biol Chem 2002, 277(18):15890–15896.

15. Goto M, Omi R, Miyahara I, Sugahara M, Hirotsu K: Structures of argininosuccinate synthetase in enzyme-ATP substrates and enzyme-AMP product forms: stereochemistry of the catalytic reaction. J Biol Chem 2003, 278(25):22964–22971.

16. Bork P, Koonin EV: A P-loop-like motif in a widespread ATP pyrophosphatase domain: implications for the evolution of sequence motifs and enzyme activity. Proteins 1994, 20(4):347–355.

17. Landrum MJ, Chitipiralla S, Kaur K, Brown G, Chen C, Hart J, Hoffman D, Jang W, Liu C, Maddipatla Z et al: ClinVar: updates to support classifications of both germline and somatic variants. Nucleic Acids Res 2025, 53(D1):D1313–D1321. Online ASS1 ClinVar database accessed 25 April 2025.

18. Brnich SE, Abou Tayoun AN, Couch FJ, Cutting GR, Greenblatt MS, Heinen CD, Kanavy DM, Luo X, McNulty SM, Starita LM et al: Recommendations for application of the functional evidence PS3/BS3 criterion using the ACMG/AMP sequence variant interpretation framework. Genome Med 2019, 12(1):3.

19. PrimateAI-3D database [https://primateai3d.basespace.illumina.com/] Accessed 7 July 2025.

20. Gao H, Hamp T, Ede J, Schraiber JG, McRae J, Singer-Berk M, Yang Y, Dietrich A, Fiziev P, Kuderna L et al: The landscape of tolerated genetic variation in humans and primates. bioRxiv 2023.

21. Woo HI, Ki CS, Lee SY, Kim JW, Song J, Jin DK, Park WS, Lee DH, Lee YW, Park HD: Mutation spectrum of the ASS1 gene in Korean patients with citrullinemia type I. Clin Biochem 2013, 46(3):209–213.

22. Martin-Hernandez E, Aldamiz-Echevarria L, Castejon-Ponce E, Pedron-Giner C, Couce ML, Serrano-Nieto J, Pintos-Morell G, Belanger-Quintana A, Martinez-Pardo M, Garcia-Silva MT et al: Urea cycle disorders in Spain: an observational, cross-sectional and multicentric study of 104 cases. Orphanet J Rare Dis 2014, 9:187.

23. Gao HZ, Kobayashi K, Tabata A, Tsuge H, Iijima M, Yasuda T, Kalkanoglu HS, Dursun A, Tokatli A, Coskun T et al: Identification of 16 novel mutations in the argininosuccinate synthetase gene and genotype-phenotype correlation in 38 classical citrullinemia patients. Hum Mutat 2003, 22(1):24–34.

24. Ruitenbeek W, Kobayashi K, Iijima M, Smeitink JA, Engelke UF, De Abreu RA, Kwast HT, Saheki T, Boelen CA, De Jong JG et al: Moderate citrullinaemia without hyperammonaemia in a child with mutated and deficient argininosuccinate synthetase. Ann Clin Biochem 2003, 40(Pt 1):102–107.

25. Haberle J, Pauli S, Schmidt E, Schulze-Eilfing B, Berning C, Koch HG: Mild citrullinemia in Caucasians is an allelic variant of argininosuccinate synthetase deficiency (citrullinemia type 1). Mol Genet Metab 2003, 80(3):302–306.

26. Moarefian S, Zamani M, Rahmanifar A, Behnam B, Zaman T: Clinical, laboratory data and outcomes of 17 Iranian citrullinemia type 1 patients: Identification of five novel ASS1 gene mutations. JIMD Rep 2022, 63(3):231–239.

27. Dimmock DP, Trapane P, Feigenbaum A, Keegan CE, Cederbaum S, Gibson J, Gambello MJ, Vaux K, Ward P, Rice GM et al: The role of molecular testing and enzyme analysis in the management of hypomorphic citrullinemia. Am J Med Genet A 2008, 146A(22):2885–2890.

28. Engel K, Hohne W, Haberle J: Mutations and polymorphisms in the human argininosuccinate synthetase (ASS1) gene. Hum Mutat 2009, 30(3):300–307.

29. Daou M, Souaid M, Yammine T, Khneisser I, Mansour H, Salem N, Nemr A, Awwad J, Moukarzel A, Farra C: Analysis of ASS1 gene in ten unrelated middle eastern families with citrullinemia type 1 identifies rare and novel variants. Mol Genet Genomic Med 2023, 11(2):e2058.

30. Barends M, Pitt J, Morrissy S, Tzanakos N, Boneh A, Newborn Screening Laboratory S: Biochemical and molecular characteristics of patients with organic acidaemias and urea cycle disorders identified through newborn screening. Mol Genet Metab 2014, 113(1-2):46–52.

31. Sander J, Janzen N, Sander S, Steuerwald U, Das AM, Scholl S, Trefz FK, Koch HG, Haberle J, Korall H et al: Neonatal screening for citrullinaemia. Eur J Pediatr 2003, 162(6):417–420.

32. Xie MJ, Cromie GA, Owens K, Timour MS, Tang M, Kutz JN, El-Hattab AW, McLaughlin RN, Jr., Dudley AM: Constructing and interpreting a large-scale variant effect map for an ultrarare disease gene: Comprehensive prediction of the functional impact of PSAT1 genotypes. PLoS Genet 2023, 19(10):e1010972.

33. Crick FH, Orgel LE: The Theory of Inter-Allelic Complementation. J Mol Biol 1964, 8:161–165.

34. ClinicalTrials.gov [https://www.clinicaltrials.gov/study/NCT06255782] OTC-HOPE trial: An Open-label Study to Investigate ECUR-506 in Male Babies Less Than 9 Months of Age With Neonatal Onset OTC Deficiency. Accessed 7 July 2025.

35. Sriretnakumar V, Harripaul R, Kennedy JL, So J: When rare meets common: Treatable genetic diseases are enriched in the general psychiatric population. Am J Med Genet A 2024, 194(8):e63609.

36. Haberle J, Vilaseca MA, Meli C, Rigoldi M, Jara F, Vecchio I, Capra C, Parini R: First manifestation of citrullinemia type I as differential diagnosis to postpartum psychosis in the puerperal period. Eur J Obstet Gynecol Reprod Biol 2010, 149(2):228–229.

37. Ziegler A, Koval-Burt C, Kay DM, Suchy SF, Begtrup A, Langley KG, Hernan R, Amendola LM, Boyd BM, Bradley J et al: Expanded Newborn Screening Using Genome Sequencing for Early Actionable Conditions. JAMA 2025, 333(3):232–240.

38. Leblond M, Galati M, Roberts J, Etheredge H, Willacy N, Ozkurt O, Pichini A: Co-Creating the Experience of Consent for Newborn Genome Sequencing: The Generation Study. Public Health Genomics 2024, 27(1):210–227.

